# Inhibition of potassium ion channels reduces Semliki Forest virus genome replication

**DOI:** 10.1101/2023.05.24.541985

**Authors:** Tristan Russell, Caoimhe O’Brien, Disha Gangotia, Stefanie Fulford, Roísín Kenny, Abdullah Alkhamees, Shonnette Premchand-Branker, Rennos Fragkoudis, Gerald Barry

## Abstract

**Introduction:** *Semliki forest virus* (SFV) is a model virus used to investigate the Alphavirus genus, which includes human pathogens Chikungunya virus and Ross River virus. Viruses harness cellular machinery to facilitate various steps of their replicative cycles. Ion channels are one group of cellular proteins required for the efficient replication of some viruses, including Influenza A viruses, Ebola virus and members of the *Betacoronavirus* genus. This study focussed on understanding SFV’s requirement for functional ion channels during replication.

**Methods:** The effect of ion channel inhibitors on *in vitro* SFV infections was measured to investigate the contribution of ion channels in its replication cycle.

**Results:** *In vitro* SFV infections carried out in the presence or absence of different ion channel inhibitors showed broad-range K^+^ channel inhibitors reproducibly attenuated virus replication and reduced its cytotoxicity in two mammalian cell lines. These broad-range K^+^ channel inhibitors disrupted an early, post-entry step causing a delay or reduction in SFV protein and RNA synthesis. Screens using inhibitors of specific K^+^ channel families showed that two-pore domain K^+^ channel (2pK) inhibitors attenuated SFV replication. Confocal microscopy revealed decreased detection of dsRNA and SFV protein in the presence of inhibitor but no change in RNA and protein colocalisation, which would indicate disruption of replication complexes. Broad-range K^+^ and 2pK inhibitors decreased viral RNA replication and transcription from the subgenomic promoter.

**Conclusions:** K^+^ channel inhibitors attenuate *in vitro* SFV replication by inhibiting an early, post-entry step of virus replication, potentially RNA synthesis.

**Importance:** No antiviral therapies have been approved for clinical use against diseases caused by members of the Alphavirus genus. Work presented in this manuscript shows for the first time that SFV genome replication and virus induced cytotoxicity can be reduced *in vitro* by treating infected cells with K^+^ channel inhibitors. This work provides the basis for investigating the effectiveness of K^+^ channel inhibitors against other alphaviruses both *in vitro and in vivo* and, because many ion channel inhibiting drugs are already in clinical use, rapid repurposing against alphavirus infections would be possible.

## Introduction

Many emerging and established pathogens of humans and animals belong to the *Alphavirus* genus including Chikungunya virus (CHIKV), which can cause a flu-like illness, rash, arthralgia, and arthritis over an extended period. Since 2006, CHIKV has spread across many parts of the world infecting thousands of people each year [1–3]. Up to the 17^th^ of March, the ECDC have reported 114,181 cases of CHIKV resulting in 43 deaths during 2023 [4, 5]. Most alphaviruses are arboviruses including CHIKV, which is transmitted by mosquitos species *Aedes aegypti* and *Aedes albopictus* [1].

As well as CHIKV, other clinically important alphaviruses include Eastern equine encephalitis virus, Western equine encephalitis virus, Venezuelan equine encephalitis virus and Ross River virus. Ross River virus is also transmitted by mosquito bites and presents in a similar way to CHIKV with fever, polyarthralgia, myalgia and a rash being typical. The equine encephalitis viruses, in contrast, are associated with neurological signs as the virus predominantly targets the brain [6–8]. In the USA between 2012 and 2021 there were 111 confirmed cases of Eastern equine encephalitis virus infection in humans causing 46 deaths while hundreds of horses were also infected [9]. In Australia there are approximately 5000-6000 confirmed human cases of Ross River virus infections per annum according to the Australian National Notifiable Diseases Surveillance System. Despite this group of viruses causing a major disease burden in humans and animals, there are no antivirals on the market. Vaccines have only recently become available for CHIKV, and the only other Alphavirus vaccines are against some equine encephalitis viruses, but they are only licensed for use in horses and need to be administered twice a year to maintain protection against severe disease.

*Semliki Forest virus* (SFV) is a model *Alphavirus*, which has been used extensively to investigate *Alphavirus* replication and pathogenesis [10–13]. SFV and other members of the *Alphavirus* genus require host factors to efficiently carry out some steps of their replication including genome replication. Genome replication takes place in spherules, which are invaginations in the membrane of large cytoplasmic vacuoles, where viral and cellular proteins required for viral RNA synthesis accumulate to form replication complexes (RCs). Viral non-structural protein 1 (nsP1) mediates localisation of viral genomes and the non-structural polyprotein (nsP1234) to the plasma membrane where the endocytic pathway is hijacked to form large cytoplasmic vacuoles [14–19]. The nsP3 proteins of SFV and another model Alphavirus, Sindbis virus, function as hub proteins at RCs because they interact with several host proteins such as the RNA-binding G3BP proteins, which contribute to viral RNA synthesis [14, 20–26].

In the cytoplasm, structural glycoproteins traffic through the golgi and endoplasmic reticulum to the plasma membrane, while capsid assembles in association with newly replicated copies of the virus genome [14, 15, 19, 27]. During its replication cycle, SFV produces a 6 kilodalton protein called 6K, which is a viroporin involved in the budding of the virus from infected cells. Mutant versions of SFV lacking 6K have normal protein production, but the deletion mutants have a defect in their assembly and budding of new particles. A recent study with Sindbis virus suggests that 6K plays a role in glycoprotein transport to the plasma membrane as well as virus particle budding from infected cells but is not involved in early stages of genome replication or protein production [28–30].

Aside from 6K viroporin, cellular ion channels also contribute to *Alphavirus* replication cycles. Cellular ion channels regulate the movement of calcium (Ca^2+^), chloride (Cl^-^), potassium (K^+^) and sodium (Na^+^) ions across membranes to maintain or change membrane potentials, which is critical to transmission of neuronal signals, muscle contractions and regulation of multiple cellular processes. Efficient CHIKV genome replication has been shown to require Cl^-^ intracellular channels 1 and 4, which interact with nsP3 and appear to contribute to RNA replication at RCs [31]. Also, inhibition of the non-selective Ca^2+^ channel, transient receptor potential vanilloid 1, which normally regulates inflammatory responses, attenuates CHIKV infection of a macrophage cell line suggesting the ion channel contributes to viral replication [32]. Ion channels are also required for the efficient replication of several other viruses [33–35]. For example, Coronaviruses and Ebola viruses both require 2-pore channel activity for efficient entry into cells via the endocytic pathway because the Ca^2+^ transport into endosomes facilitates fusion of the viral envelope with the endosomal membrane and release of the genome into the cytoplasm [36–39]. During the replication of some *Peribunyaviridae*, the influx of K^+^ into virus-containing endosomes through K^+^ channels causes conformational changes in their envelope proteins, which also facilitates release of their genomes into the cytoplasm [40, 41]. Beyond their *in vitro* and *in vivo* antiviral effects in laboratory settings, some ion channel inhibitors have been used to treat viral diseases in humans [42–45].

The role of ion channels in SFV replication was investigated by carrying out *in vitro* infections of mammalian cell lines in the presence or absence of ion channel inhibitors targeting different ion channels and measuring their impact on virus replication and cytotoxicity. K^+^ channel inhibitors reproducibly caused a significant reduction in SFV replication and cytotoxicity. K^+^ channel inhibitors were not effective at high MOI suggesting they are not potent but were effective when administered after SFV infections were already established or when administered to SFV-infected mosquito cells. Mechanistic studies showed K^+^ channel inhibitors disrupt an early, post-entry step, which confocal imaging and a transreplication assay revealed was most likely viral RNA synthesis. This work has shown K^+^ channels are required for efficient *in vitro* replication of SFV, which provides grounds to assess the ability of K^+^ channel inhibitors to attenuate *in vivo Alphavirus* infections or infections caused by other, pathogenic *Alphavirus* species.

## Methods

### Cell lines

NIH 3T3 mouse and Baby Hamster Kidney-21 (BHK-21) fibroblast cells were cultured in Dulbecco’s Modified Eagle Medium (DMEM) supplemented with 10% new-born calf serum or 5% foetal bovine serum, respectively. Cells were grown in incubators held at 37°C and in a 5% CO₂ atmosphere. *A. aegypti* Aag.2 cells were maintained in Drosophila media supplemented with 10% foetal bovine serum and were incubated at 28°C.

### Virus

SFV was generated using a reverse genetics approach [46]. BHK-21 cells grown to approximately 90% confluency in T-175 tissue culture flasks were transfected with a plasmid containing the full-length wild type SFV4 genome (pCMV-SFV4) using Lipofectamine 2000 following the manufacturer’s protocol with some modifications. Old growth media was replaced with 20 ml fresh growth media and a solution containing 2% lipofectamine 2000 reagent and 1 μg pCMV-SFV4 was made up and added to the tissue culture flask. Cells were monitored microscopically for signs of cytopathic effects (CPE) indicative of virus infection. Two or three days post transfection, when at least 90% of cells displayed CPE, virus was harvested.

### Plaque assays

Plaque assays were used to obtain SFV titres including stocks. BHK-21 cells grown in 6-well plates until approximately 70-80% confluent were infected with serially diluted SFV. SFV was adsorbed for 120 minutes then aspirated from the cells before addition of growth media containing 1.25% Avicel (FMC Biopolymer) thickening agent. Infected cells were returned to the incubator for 3 days, at which point, cells were fixed in 4% paraformaldehyde (PFA) and stained with 5% crystal violet. Plaques were counted and the PFU/ml calculated.

### SFV infections and ion channel inhibitor treatments

SFV infections were carried out at the specified multiplicity of infection (MOI). Growth media was removed from cells when approximately 80-90% confluent, SFV stock was diluted in an appropriate volume of growth media and added to cells, which were returned to the incubator for 2 hours adsorption. For infections at high MOI (> 0.1), virus was removed, cells were washed twice in PBS to remove extracellular virus and then fresh growth media was added to cells. Unless otherwise stated, PBS washes were not carried out for infections at low MOI (0.001).

Cells were treated with ion channel inhibitors at concentrations shown to be non-cytotoxic (Table 1). For all inhibitors used, untreated controls were set up by carrying out infections in the presence of the solvent used to make inhibitor solutions (Table 1). Ion channel inhibitors were added at the specified time relative to infection. When inhibitors were added at 0 hours post-infection (hpi), adsorptions were carried out in the presence of the inhibitor for 2 hours, then replaced with fresh growth media containing inhibitor at the working concentration. When added at later time points (at least 4 hpi), inhibitor was added directly to the media to achieve the working concentration without removing the existing growth media.

**Table 1:** Inhibitors used in drug screens. All inhibitors were manufactured and supplied by Sigma Aldrich, except Tram-34 (Tocris Bioscience).

| Inhibitor | Solvent | Working concentration | Ion channel target |
| --- | --- | --- | --- |
| 2-aminoethyl diphenylborinate (2-APB) | DMSO | 10 μM | Ca<sup>2+</sup> |
| 4-aminopyridine (4-AP) | Water | 1 mM | Voltage-gated K<sup>+</sup> |
| Clotrimazole | DMSO | 1 μM | K<sup>+</sup> |
| 4,4'-Diisothiocyanatostilbene-2,2'-disulfonic acid disodium salt hydrate (DIDS) | DMSO | 10 μM | Cl<sup>-</sup> |
| Disopyramide phosphate (DPO) | Water | 90 μM | Na<sup>+</sup> |
| Glyburide | DMSO | 20 μM | ATP-dependent K<sup>+</sup> |
| Indanyloxyacetic acid 94 (IAA-94) | DMSO | 100 μM | Cl<sup>-</sup> |
| Lidocaine | Ethanol | 200 μM | Na<sup>+</sup> |
| Potassium chloride (KCl) | Water | 50 mM | K<sup>+</sup> |
| Ruthenium red | Water | 50 μM | Two-pore K<sup>+</sup> |
| Spermine | Water | 250 μM | Two-pore K<sup>+</sup> |
| Tetraethylammonium chloride (TEA-Cl) | Ethanol | 8 mM | K<sup>+</sup> |
| Tram-34 | DMSO | 3 μM | Ca<sup>2+</sup>-activated K<sup>+</sup> |
| Verapamil hydrochloride | Ethanol | 10 μM | Ca<sup>2+</sup> |

### Growth curves

Growth curves were generated by infecting and treating cells grown in 24-well plates. To measure the effect of inhibitor treatments on virus replication, 100 μl of media containing the virus was collected at the specified timepoint. The effects of inhibitors on SFV titre overtime were determined by measuring the amounts of virus in collections using Tissue culture infectious dose 50 (TCID50) assays. TCID50s were set up by infecting BHK-21 cells grown in 96-well plates with serial dilutions of collected virus in quadruplicate. Wells were scored for the presence or absence of CPE three days post-infection. The Reed-Meunch TCID50 calculator was used to estimate virus titres.

### Cell viability assay

The Wst-1 cell viability assay was used to identify ion channel inhibitors that reduced SFV cytotoxicity. Cells grown in 96-well plates were infected with SFV (MOI 0.1) or mock infected in the presence or absence of ion channel inhibitors. For infections of 96-well plates, virus containing media was not replaced after two hours to reduce the risk of disrupting the cell monolayer. At 22 hpi, virus containing media was aspirated from cells and then 100 μl fresh growth media plus 10 μl of Wst-1 reagent was added to each well. Plates were returned to the incubator for 2 hours to allow metabolically active cells to convert the Wst-1 reagent to a formazan dye. Spectrophotometric readings at 450 nm were taken using the Promega Glomax Multi Detection System microplate reader at 24 hpi. Higher spectrophotometric readings were indicative of increased cell viability.

### Insect cell line infections

Aag.2 mosquito cells transfected with plasmid encoding Firefly luciferase (FLuc) and cultured for 24 hours in the presence of different concentrations of KCl. FLuc production was measured to identify non-cytotoxic concentrations of KCl. Cells were infected with recombinant SFV expressing FLuc at the indicated MOI in the presence of the KCl at concentrations shown to be non-cytotoxic. Replication of recombinant SFV produces FLuc protein so at the indicated timepoints, FLuc substrate was added to cells and luminescence readings were taken.

### Western blot

Protein was isolated from cells at the stated timepoint using lysis buffer (1 % Triton-X, 25 mM HEPES (pH 7.5), 150 mM NaCl). Growth media was removed, cells were washed once in ice-cold PBS then incubated in ice-cold lysis buffer at 4°C for 20 minutes with agitation every 4-5 minutes. Cells were detached from the plate surface using a cell scraper to aid lysis. Lysates were cleared to remove precipitated cell debris by centrifugation at 20,000 x g for 7 minutes at 4°C. Proteins in lysates were separated by SDS-PAGE and analysed by western blot. Blots were blocked in in 5% bovine serum albumin (BSA). Blots were incubated in primary antibody wash solutions (made up in 5% BSA) containing 1:3000 dilutions of rabbit anti-nsP3 antibody and 1:5000 dilutions of mouse anti-β-actin antibody at 4°C for 16-18 hours. Secondary antibody wash solutions containing 1:5000 dilutions of horseradish peroxidase conjugated anti-rabbit and anti-mouse antibodies were incubated with blots for 180 minutes at room temperature. The SuperSignal West Pico PLUS Chemiluminescent Substrate kit (Thermo Fisher Scientific) and SynGene gel imager were used to visualise antibodies bound to blots.

Protein band intensities were measured using ImageJ and nsP3 band intensities were normalised against β-actin band intensities.

### Immunofluorescence microscopy

Cells grown on coverslips in 6-well plates until approximately 70% confluency were infected with SFV or transfected with pCMV-SFV4 in the presence or absence of inhibitor. Cells were fixed in 4% PFA at the stated timepoints and then underwent fluorescent staining. Fixed cells were permeabilised by incubating in 0.3% Triton-X for 20 minutes and cells were blocked in 5% BSA. Cells were incubated in primary antibody at 37°C for 180 minutes or at 4°C for 16-18 hours. Primary antibody washes contained rabbit anti-nsP3 (1:3000) and/or mouse J2 anti-dsRNA antibody (1:500). Cells were incubated in secondary antibody wash solutions for 120 minutes at room temperature. Antibodies conjugated to AlexaFluor-488 or –594 (1:3000) were used for secondary antibody washes. Cell nuclei were stained using 1:1000 dilutions of DAPI. Cells fixed to coverslips were mounted onto microscope slides using the VectaShield HardSet mounting media. Conventional microscopy images were taken using Nikon Eclipse E400 IF microscope and images were manipulated and merged using ImageJ. Confocal images were also taken using the Zeiss 510UV Meta microscope and image analysis was carried out using both the Autoquant X3 analysis software and ImageJ. Multiple images were taken, and representative images are shown in the figures.

### Quantitative reverse-transcription PCR

BHK-21 cells grown in 24-well plates were infected with SFV (MOI 0.001) or mock infected in the presence or absence of KCl in triplicate. Cells grown in 24-well plates were also transfected with RNA encoding the SFV non-structural proteins conjugated to ZSgreen in the presence or absence of KCl. RNA was isolated at 3 hpi from infected cells and at 5, 6 and 7 hours post-transfection (hpt) from transfected cells using the Qiagen RNeasy kit as described in the manufacturer’s protocol. Cells were lysed in 350 μl of Buffer RLT and RNA was quantified using the nanodrop. RNA was diluted in nuclease-free water, so each sample contained 500 ng RNA in a final volume of 10 μl. RNA was converted to cDNA using the High Capacity cDNA Reverse Transcription kit (Life Technologies). Reaction mixes containing reverse transcriptase, random primers and nucleotides were made up as described in the manufacturer’s protocol and then incubated at 37°C for 120 minutes. Enzyme was deactivated by heating to 95°C for 5 minutes. For each condition, reverse transcriptase was replaced with nuclease free water in the cDNA synthesis step of one sample, which was the control to detect genomic DNA.

RT-qPCR of cDNA samples was carried out in duplicate using primers overlapping the region of the non-structural polyprotein that encodes nsP3 (Table 2). Positive control primers amplifying the PPIA gene of the Syrian Golden hamster were included to standardise samples (Table 2). RT-qPCR was carried out using the PowerUp SYBR green master mix kit. To make up reaction mixes, cDNA was diluted 1:4 using EDTA buffer containing 2.5 ng/μl lambda DNA and 5 μl of these dilutions was combined with 300 nM of forward and reverse primers plus 10 μl of the 2X PowerUp master mix in a final volume of 20 μl. Standard curves and no template controls (NTCs) were set up by replacing the 5 μl cDNA with 5 μl standards or dilution buffer. A nsP3 standard curve was generated by making serial 1:10 dilutions of linearised pCMV-SFV4 and carrying out RT-qPCR on these samples containing known copy numbers. A standard curve was generated for the PPIA primer pair by pooling 5 μl of each cDNA sample and making serial 1:4 dilutions of this pool. NTCs of nuclease-free water and dilution buffer (2.5 ng/μl lambda DNA) were also included. The standard PCR conditions described in the manufacturer’s protocol were used.

**Table 2:**
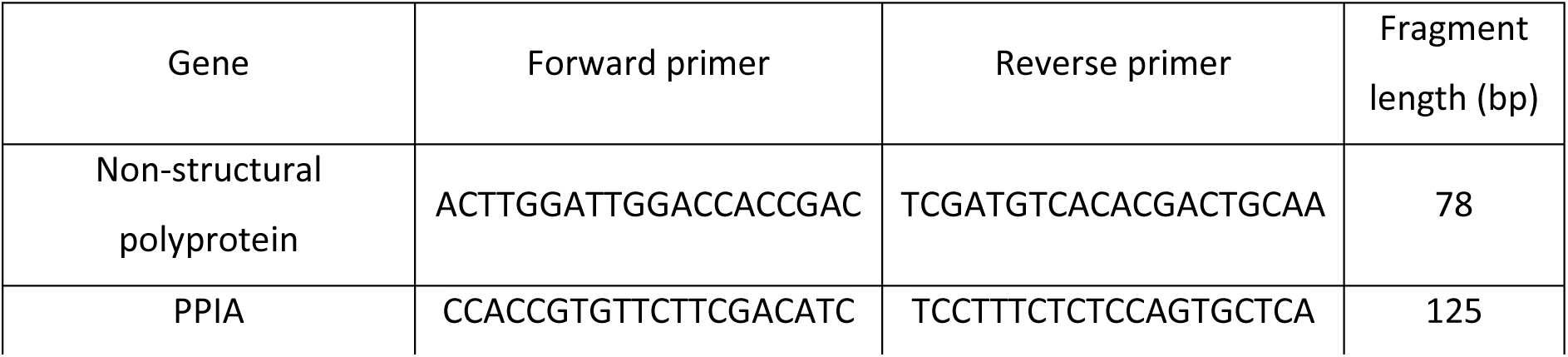
Target gene, primer sequences (5’ to 3’) and the length of the fragment generated.

A standard curve was generated for viral RNA by averaging the Ct (cycle threshold) values obtained for technical duplicates of RT-qPCR reactions of linearised pCMV-SFV4 using the viral primer pair and plotting these against the log concentration. The gradient of the slope was calculated and used to calculate the concentrations of viral RNA in samples based on Ct values, which had been standardised based on the corresponding Ct values obtained for the PPIA primer pair. Viral RNA concentrations were compared using Student t-tests as described previously.

### Transreplication assay

Transreplication assays were used to determine the effect of ion channel inhibitors on replication of genomic RNA and transcription from the sub-genomic promoter [47]. Cells were transfected with two of three plasmids in the presence or absence of ion channel inhibitors. All cells were transfected with a template plasmid (pF4-Temp) containing the Firefly luciferase (FLuc) gene downstream of the SFV 5’ untranslated region (UTR) and the Gaussia luciferase (GLuc) gene downstream of the SFV sub-genomic promoter. For test samples, cells were transfected with a plasmid encoding the wild type nsP1234 polyprotein downstream of a CMV promoter (pWTnsP1234). For control samples, cells were transfected with plasmid encoding mutant nsP1234, which cannot be cleaved at the nsP3/4 junction so no longer functions as a replicase (pMUTnsP1234). Initially wild type nsP1234 expressed in cells transfected with pWTnsP1234 would synthesise FLuc from the 5’ UTR region of pF4-Temp, then following nsP2 cis-acting protease activity, the conformation of nsP1234 would change and it would specifically synthesise GLuc from the sub-genomic promoter. At 18 hpt of BHK-21 cells, the dual-luciferase reporter assay system (Promega) was used to detect levels of FLuc and GLuc. The manufacturer’s guidelines were followed, which involved lysing the cells, and addition of FLuc then GLuc substrate to the lysate. After addition of each substrate, luminescence was read using the 20/20 Glomax Luminometer (Promega). Luminescence readings taken for test samples transfected with pWTnsP1234 were normalised against readings for control samples transfected with pMUTnsP1234.

### Statistical analysis

Ion channel inhibitors associated with significantly decreased virus titre or increased cell viability were identified by carrying out Student t-tests in R studio to compare titres obtained for treated samples and untreated controls. A p-value cut-off of < 0.05 was used to indicate a significant effect.

## Results

### Broad-range K^+^ channel inhibitors attenuate SFV replication

To investigate the role of ion channels in SFV replication, NIH 3T3 cells were infected with SFV in the presence of broad-range ion channel inhibitors targeting Cl^-^, Ca^2+^, Na^+^ or K^+^ ion channels or their solvents (untreated controls) (Table 1). Virus collected at 24, 48 and 72 hpi was measured by TCID50 (Figure 1). Ca^2+^ (Figure 1A) and Cl^-^ (Figure 1B) channel inhibitors had no significant effect on SFV titre suggesting the ion channels targeted by these inhibitors are not involved in SFV replication. One Na^+^ channel inhibitor, lidocaine, caused a significant reduction in the SFV TCID50/ml at 24 hpi, but the other Na^+^ channel inhibitor tested, Disopyramide phosphate (DPO), had no effect (Figure 1C). SFV titres were significantly reduced at 24 and 48 hpi when cells were infected in the presence of the K^+^ channel inhibitors potassium chloride (KCl) and Tetraethyl-ammonium chloride (TEA-Cl) while clotrimazole only caused a significant reduction at 48 hpi (Figure 1D). No ion channel inhibitors caused a significant reduction in SFV titre for collections made at 72 hpi (Figure 1C). Growth curve assays with drugs shown to have a significant effect in NIH 3T3 cells were repeated in BHK-21 cells to test for cell-type specific effects (Figure 1E). The attenuation of SFV replication caused by KCl and TEA-Cl in NIH 3T3 cells was reproducible in BHK-21 cells at 24 hpi, but not at 48 hpi (Figure 1E). Clotrimazole and lidocaine did not cause a significant decrease in SFV replication at any timepoint in BHK-21 cells.

**Figure 1:**
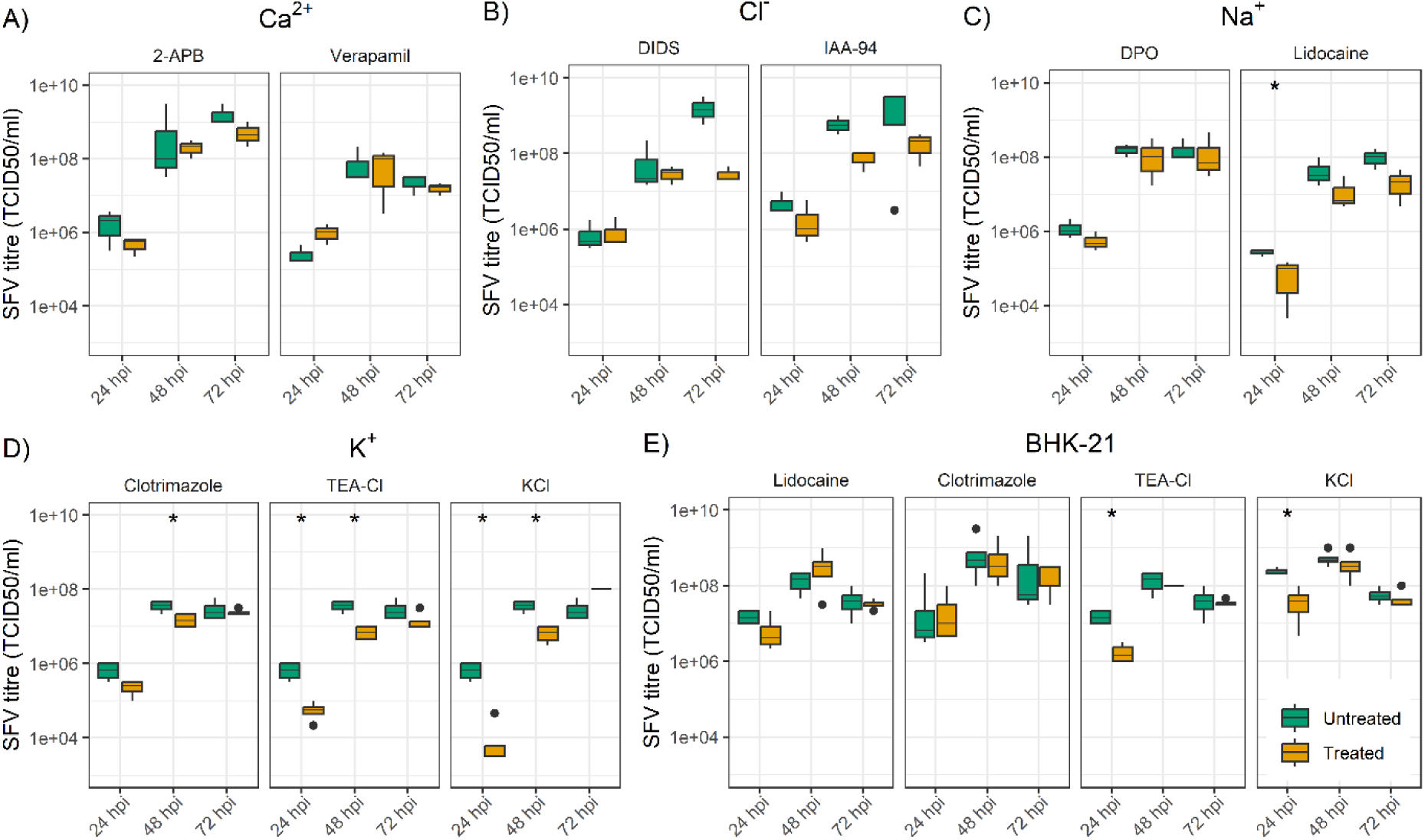
Growth curve assays with broad-range inhibitors. NIH 3T3 cells were infected with SFV (MOI 0.001) in the presence of broad-range inhibitors of Cl^-^ (A), Ca^2+^ (B), Na^+^ (C) or K^+^ (D) channels or the solvent used to make drug stocks (untreated). Media was collected at the stated timepoints and TCID50s for treated and untreated controls were compared by Student t-tests. E) Selected inhibitors were also screened in BHK-21 cells. The box of boxplots represents the upper and lower quartile, the line the median, the whiskers the range and dots the outliers of at least three repeats. “*” indicates inhibitors where the Student t-test p-value was < 0.05. Abbreviations: 2-APB, 2-aminoethoxydiphenyl borate; DIDS, 4,4’-diisothiocyano-2,2’-stilbenedisulfonic acid; DPO, diphenyl phosphine oxide 1; IAA– 94, indanyloxyacetic acid-94; KCl, potassium chloride; and TEA-Cl, tetraethylammonium chloride.

### K^+^ channel inhibitors protect SFV infected cells

Cell viability assays using broad-range ion channel inhibitors were carried out to see if virus cytotoxicity was reduced. BHK-21 cells were infected with SFV (MOI 0.1) in the presence of inhibitors or their solvents (untreated controls), then cell viability was measured using a WST-1 assay at 24 hpi (Figure 2). The Ca^2+^ channel inhibitor 2-Aminoethyl diphenylborinate (2-APB) (Figure 2A) and the Cl^-^ channel inhibitor 4,4′-Diisothio-cyanatostilbene-2,2′-disulfonic acid disodium salt hydrate (DIDS) (Figure 2B) significantly increased WST-1 absorbance readings. The second Ca^2+^ channel (verapamil hydrochloride) and Cl^-^ channel (Indanyloxyacetic acid 94) inhibitors used had no effect on absorbance readings (Figure 2A, 2B). The Na^+^ channel inhibitors also had no significant effect on cell viability (Figure 2C). A significant increase in WST-1 absorbance readings was measured when BHK-21 cells were infected with SFV in the presence of the K^+^ channel inhibitors KCl and TEA-Cl compared to untreated controls (Figure 2D).

**Figure 2:**
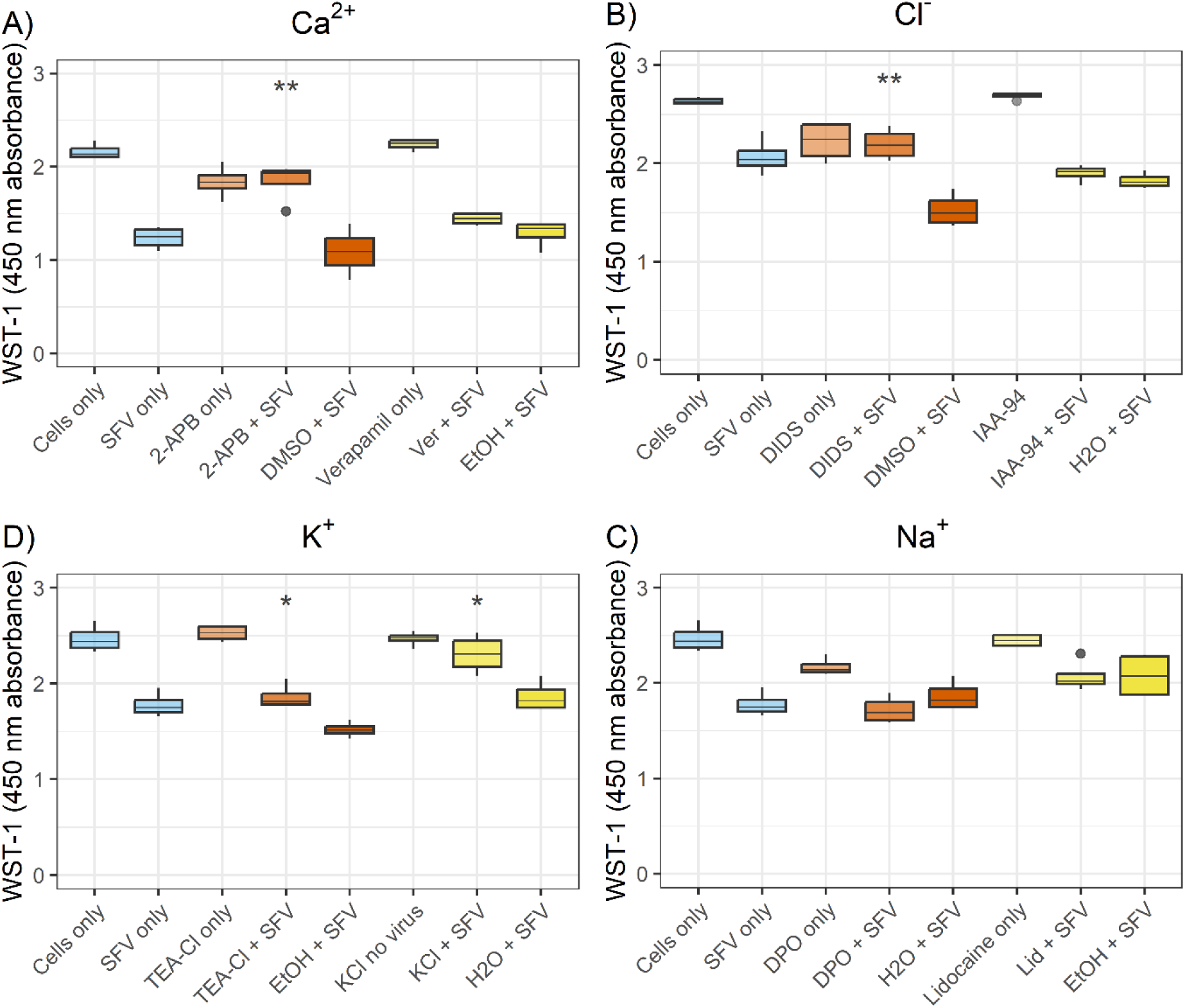
Cell viability screen of broad-range inhibitors. WST-1 was used to measure the viability of BHK-21 cells infected with SFV (MOI = 0.1) in the presence of broad-range Ca^2+^ (A), Cl^-^ (B), Na^+^ (C) or K^+^ (D) channel inhibitors or solvents used to make inhibitor stocks (untreated controls). Untreated/uninfected cells (cells only), untreated/infected cells (SFV only) and treated/uninfected cells were included as controls. WST-1 assays were performed at 24 hpi. Plots show absorbance readings for each sample with high readings indicating increased cell viability. Inhibitors and their corresponding untreated control were plotted in the same colour. Student t-tests were used to compare absorbance readings for treated samples and untreated controls. “*” indicates inhibitors with p-values of 0.005-0.05 and “**” indicates inhibitors with p-values 0.0005-0.005. Abbreviations: 2-APB, 2– aminoethoxydiphenyl borate; DIDS, 4,4’-diisothiocyano-2,2’-stilbenedisulfonic acid; DMSO, dimethylsulfoxide; DPO, diphenyl phosphine oxide 1; EtOH, ethanol; IAA-94, indanyloxyacetic acid-94; Lid, lidocaine; KCl, potassium chloride; TEA-Cl, tetraethylammonium chloride; and Ver, verapamil.

### Characterising the inhibition of SFV replication by KCl

Results of the growth curve and cell viability assays with broad spectrum inhibitors supported pursuing K^+^ channels for further investigation into their potential role in SFV replication. The WST-1 cell viability assay was repeated in NIH 3T3 cells with a second dose of 50 mM KCl added at 8 hpi after infecting cells in the presence of KCl. Treatment with a single dose or a double dose of KCl significantly increased WST-1 absorbance readings compared to untreated controls suggesting KCl protects NIH 3T3 cells from SFV-induced CPE (Figure 3A). However, there was no difference in the level of protection provided by the single and double doses (Figure 3A).

**Figure 3:**
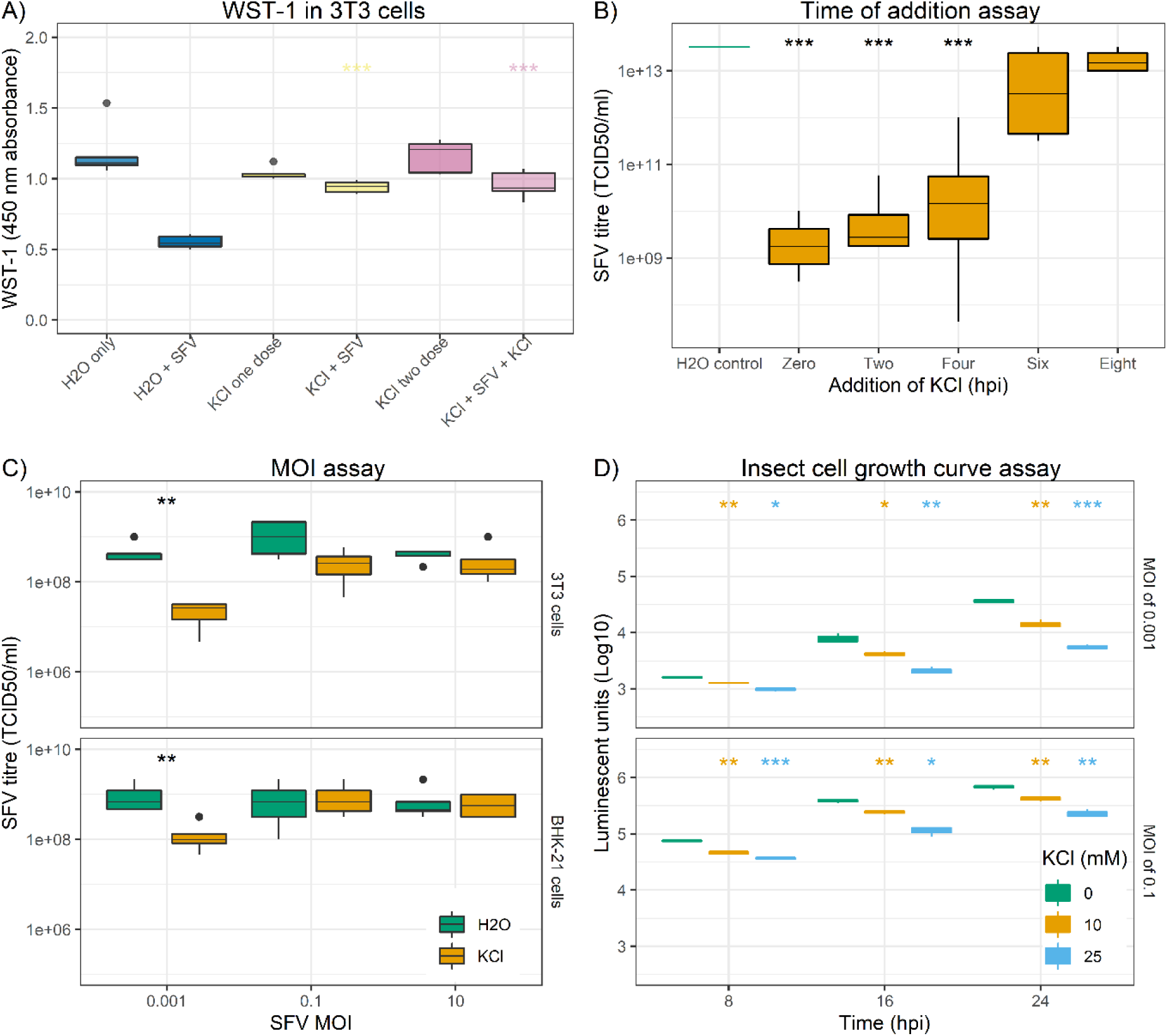
Understanding the effect of KCl on in vitro SFV replication. A) A WST-1 cell viability assay was carried out in SFV-infected NIH 3T3 cells where KCl was administered at 0 hpi (KCl + SFV) and at 0/8 hpi (KCl + SFV + KCl). B) Cells infected with SFV (MOI 0.001) were treated with 50 mM KCl at the specified timepoint post-infection. Media was collected 24 hpi and the amount of virus in samples measured by TCID50. C) Cells were infected with SFV at different MOI in the presence or absence of 50 mM KCl and the effect on virus at 24 hpi measured by TCID50. D) Aag.2 cells were infected with recombinant SFV encoding Firefly luciferase at MOI 0.001 and 0.1 in the presence of 0, 10 and 25 mM KCl and replication was measured using a luciferase assay. Significant results were labelled as described in Figure 2 and “***” indicated results with p-value <0.0005.

A time of addition assay was carried out to determine the ability of KCl to attenuate SFV replication when the infectious cycle was already underway. NIH 3T3 cells were infected with SFV and 50 mM KCl was added 0, 2, 4, 6 or 8 hpi, then virus collected at 24 hpi underwent TCID50. Compared to the untreated control, addition of KCl at each timepoint tested reduced SFV titre at 24 hpi, but the difference only reached significance when KCl was added at 0, 2 or 4 hpi (Figure 3B). The potency of KCl was tested by infecting NIH 3T3 and BHK-21 cells in its presence or absence at MOI 0.1 and 10 because a more potent inhibitor would be expected to remain effective at higher MOI (Figure 3C). Cells were also infected at MOI 0.001 as KCl has already been shown to be effective at this concentration. PBS washes were carried out after adsorptions for infections at each MOI. While KCl remained effective when infections were carried out at MOI 0.001, it did not cause any significant decrease in SFV titre at MOI 0.1 or 10 (Figure 3C).

SFV is transmitted by the *Aedes* mosquito so the effect of KCl on SFV replication in the *Aedes aegypti* cell line, Aag.2 was assessed using a luciferase assay (Figure 3D). KCl was added at 25 mM and 10 mM because cytotoxic effects were observed when 50 mM was used. There was a dose-dependent effect on SFV replication as measured by the luciferase assay in Aag.2 cells and KCl was effective when cells were infected at 0.001 and 0.1 MOI (Figure 3D).

### KCl inhibits an early step in SFV replication

The effect of KCl on SFV non-structural protein was assessed as an effect at protein abundance would indicate KCl disrupts an early step of SFV replication. SFV non-structural proteins are translated as a polyprotein following release of the viral genome into the cytoplasm [48]. Western blot and immunofluorescence microscopy (IF) with an anti-nsP3 antibody were used to detect viral protein abundance at the specified timepoints. Infecting BHK-21 cells with SFV in the presence of KCl delayed and reduced nsP3 detection by western blot with no effect on the cellular protein β-actin (Figure 4A). There was also a delay and reduction in nsP3 detection by IF when NIH 3T3 cells were infected with SFV in the presence of KCl (Figure 4B). Viral RNA synthesis also occurs early in the SFV replication cycle and it was shown treating SFV-infected BHK-21 cells with KCl caused a significant reduction in viral RNA abundance at 3 hpi (Figure 4C).

**Figure 4:**
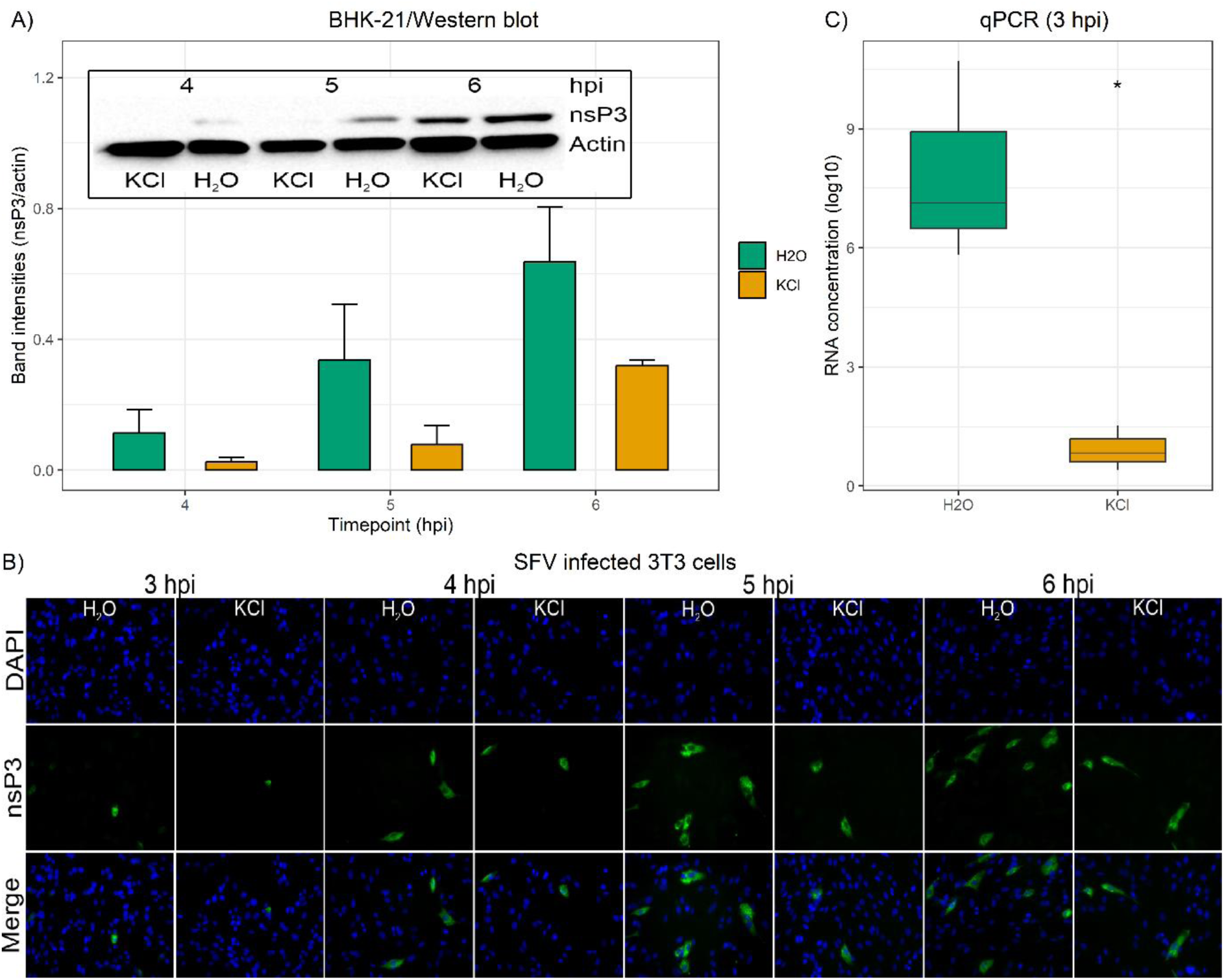
KCl inhibits an early step of SFV replication. A) BHK-21 cells were infected with SFV (MOI 0.1) in the presence or absence of KCl and lysed at the stated timepoints. Lysates were analysed by western blot using antibodies against β-actin and nsP3. Band intensities were measured using ImageJ and normalised. B) NIH 3T3 cells were infected with SFV (MOI 0.1) in the presence or absence of KCl and fixed at the stated timepoints. Fixed cells were immunostained with anti-nsP3 antibody (green). C) BHK-21 cells were infected with SFV (MOI 0.001) in the presence or absence of KCl, then RNA was isolated at 3 hpi for analysis by qPCR. Significant results were annotated as in Figure 1.

### K+ channels are involved in a post-entry step

Ion channels have previously been shown to be required for the entry of viruses including a role for K^+^ channels in the entry of Peribunyaviruses where they facilitate fusion of the viral envelope with the endosomal membrane to release the genome into the cytoplasm [40, 41, 49]. Alphaviruses also hijack the endocytic pathway to enter cells [50] so the effect of KCl on SFV entry was investigated by bypassing the entry step. Entry was bypassed by transfecting cells with pCMV-SFV4, which delivers the RNA genome to the cytoplasm via a transcription of a transfected plasmid rather than using the endocytic pathway. Cells were transfected with pCMV-SFV4 in the presence or absence of KCl and viral protein was detected. Disruption of transfection and exogenous gene expression by KCl was determined by transfecting cells with a STAT1-GFP plasmid with or without KCl, but there was no change in protein expression as measured by IF (Figure 5A).

**Figure 5:**
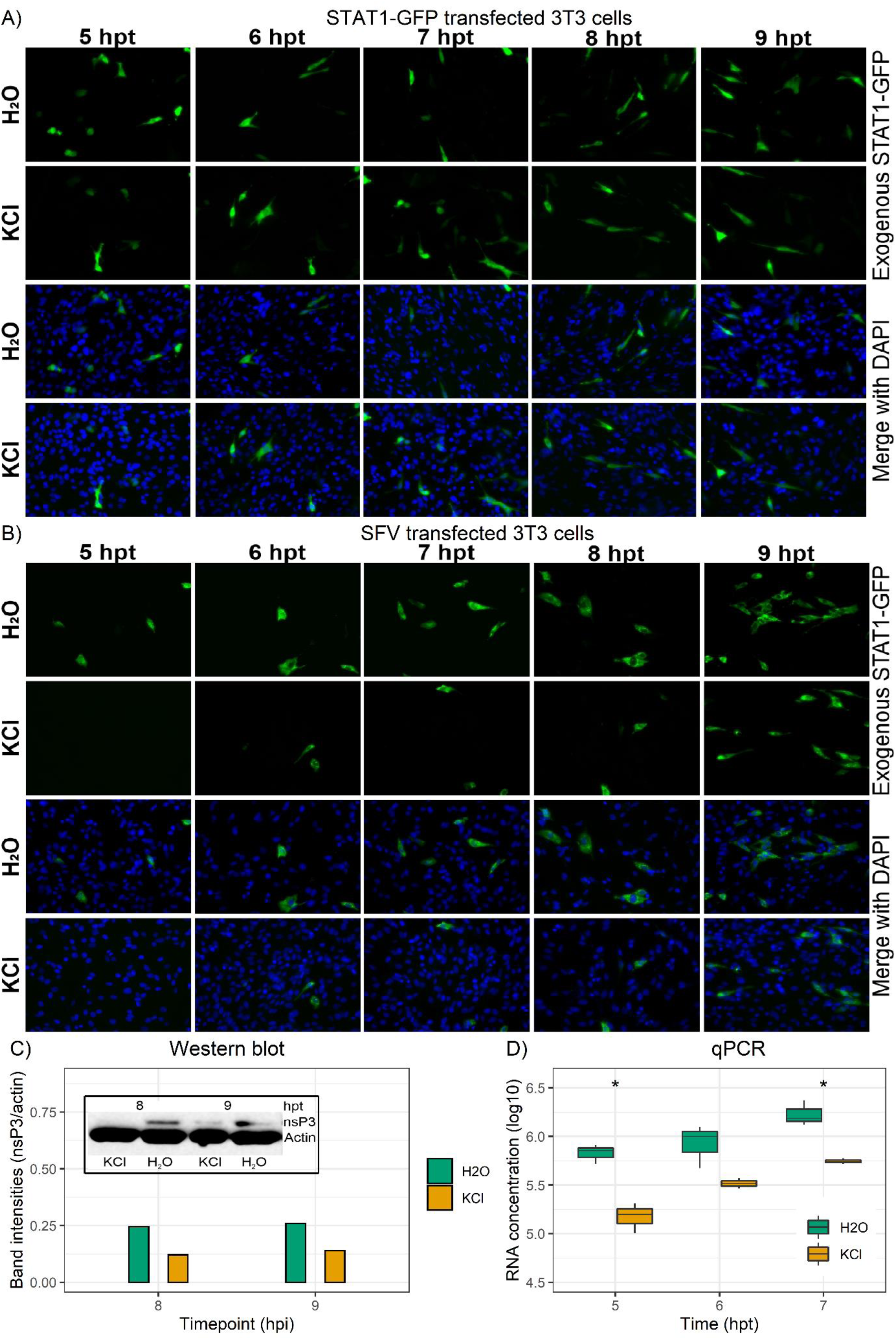
K^+^ channel inhibitors disrupt an early, post-entry step of SFV replication. A) NIH 3T3 cells grown in 6-well plates were transfected with 1 μg of plasmid encoding STAT1-GFP in the presence or absence of KCl and fixed at the specified timepoints. B) NIH 3T3 cells grown in a 6-well plate were transfected with 1 μg of pCMV-SFV4 in the presence or absence of KCl. Cells were then fixed at the stated timepoints and immunostained with anti-nsP3 antibody (green). C) BHK-21 cells grown in 6-well plates were transfected with 1 μg of pCMV-SFV4 and lysed at the specified timepoints. Lysates were then analysed by western blot using antibodies against β-actin and nsP3. Band intensities were measured using ImageJ and normalised. D) RNA abundance as measured by qPCR for BHK-21 cells transfected with 1µg RNA encoding SFV non-structural proteins in the presence or absence of KCl. Significant results were annotated as in Figure 1.

There was a reduction in the number of NIH 3T3 cells where nsP3 was detectable by IF following pCMV-SFV4 transfection in the presence of KCl (Figure 5B). At 5 hours post-transfection (hpt) there was no nsP3 detected in the KCl-treated cells but nsP3 was detected in the untreated cells (Figure 5B). The same delay and reduction were observed when BHK-21 cells were transfected with pCMV-SFV4 in the presence of KCl and nsP3 was detected by western blot (Figure 5C). A decrease in viral RNA abundance was also seen when BHK-21 cells were transfected with RNA encoding the non-structural proteins conjugated to ZSgreen in the presence of KCl, which delivers the RNA directly to the cytoplasm (Figure 5D). This shows KCl disrupts viral RNA synthesis by the non-structural proteins expressed from this RNA and indicates the effect of KCl on protein abundance in the cells transfected with pCMV-SFV4 is not just due to inhibition of plasmid transcription.

### Two-pore domain inhibitors disrupt SFV replication

Screens with family-specific K^+^ channel inhibitors were carried out to narrow down the types of K^+^ channel involved in SFV replication (Figure 6). The family-specific K^+^ channel inhibitors used were Tram-34, glyburide, 4-aminopyridine (4-AP) and ruthenium red, which target Ca^2+^-activated, ATP-dependent, voltage-gated and 2-pore domain K^+^ channels (2pK), respectively [51–56]. Tram-34, glyburide and 4-AP had no significant effect on SFV titres indicating the K^+^ channels targeted by these inhibitors are not involved in SFV replication in NIH 3T3 cells (Figure 6A). Treating SFV-infected NIH 3T3 cells with the 2pK inhibitor ruthenium red significantly reduced SFV TCID50 and a second 2pK inhibitor, spermine, also caused a reduction at 24 hpi (Figure 6A). Both 2pK inhibitors also caused a significant reduction in SFV titre when the assay was repeated in BHK-21 cells using either TCID50 (Figure 6B) or plaque assay (Figure 6C) to titrate virus.

**Figure 6:**
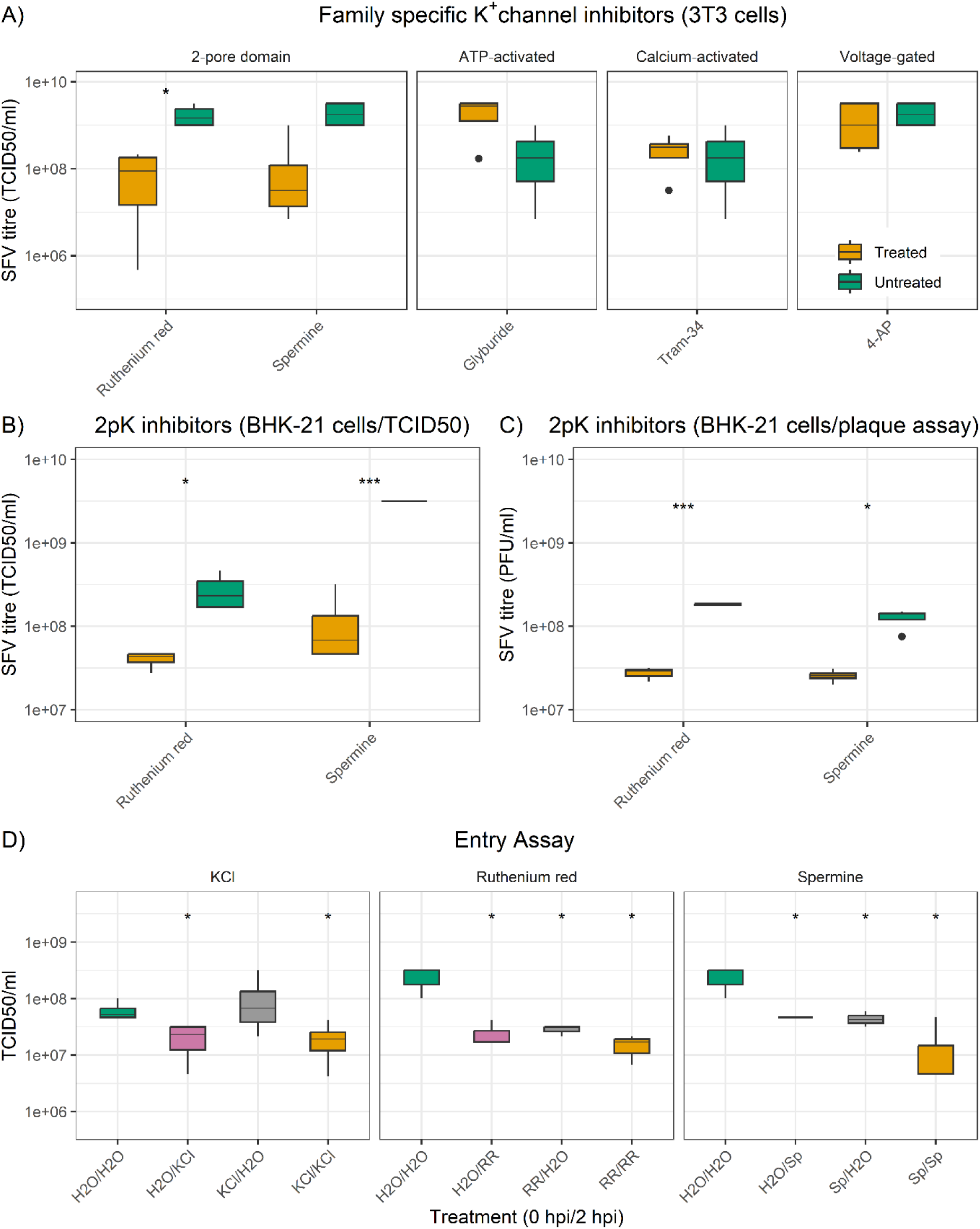
2pK inhibitors are the only family specific K^+^ channel inhibitors that attenuate SFV replication. A) NIH 3T3 cells were infected in the presence of family-specific K^+^ channel inhibitors, then virus collected at 24 hpi was titrated by TCID50. B) The assay in A) was repeated in BHK-21 cells. C) The assay in A) was repeated in BHK-21 cells but using plaque assay to titrate virus. D) BHK-21 cells were infected with SFV (MOI 0.001) in the presence or absence of inhibitors at different stages of the infection relative to the 2-hour adsorption. Virus collected at 24 hpi was titrated by TCID50. Significance was annotated as in previous figures. Abbreviations: RR, ruthenium red; Sp, spermine.

The effect of 2pK inhibitors, as well as KCl, on SFV entry was assessed by adding inhibitors during the 2-hour adsorption, but not post-adsorption, or only adding inhibitors post-adsorption so they were present or absent during the entry step, respectively. KCl caused a significant reduction in SFV titre when added post-adsorption but not when it was only present during the adsorption period providing additional evidence that KCl disrupts a post-entry step (Figure 6C). Ruthenium red and spermine both caused a significant decrease in SFV titre when added at each time relative to adsorption (Figure 6C). This could suggest they disrupt SFV entry, or it could indicate a more lasting effect on K^+^ channel activity when compared to KCl.

### K^+^ channel inhibitors attenuate viral RNA synthesis

Previous results have shown KCl disrupts an early, post-entry step of SFV replication as supported by the delay and reduction in SFV nsP and RNA abundance even when entry was bypassed. As the same cellular machinery required for translation of the nsP1234 polyprotein is also used to translate cellular proteins, such as β-actin, and non-viral exogenous proteins, such as STAT1-GFP, for which no effect of KCl was found, it suggests KCl is disrupting viral RNA synthesis. Synchronised SFV infections of BHK-21 in the presence or absence of KCl were co-immunostained for nsP3 and dsRNA to simultaneously assess the effect of KCl on abundance of nsP and viral RNA. Confocal images of cells fixed at 4 hpi appeared to show a decrease in the abundance of nsP3 and dsRNA staining (Figure 7A). Comparing intensity measurements for treated and untreated showed there was a significant decrease in the level of staining for both viral components (Figure 7B). BHK-21 cells were also transfected with pCMV-SFV4 in the presence or absence of KCl and fixed at 0.5 hours intervals from 5 to 9 hpt. As with the synchronised infections, there appeared to be a decrease in staining of viral components in treated cells with this difference most obvious at the earlier timepoints, and in general, the difference in dsRNA staining appeared greater than the difference in nsP3 staining (Figure 8A). This was supported by the intensity readings, which showed there was a significant difference in dsRNA staining between treated and untreated, but there was no statistically significant difference in nsP3 staining (Figure 8B).

**Figure 7:**
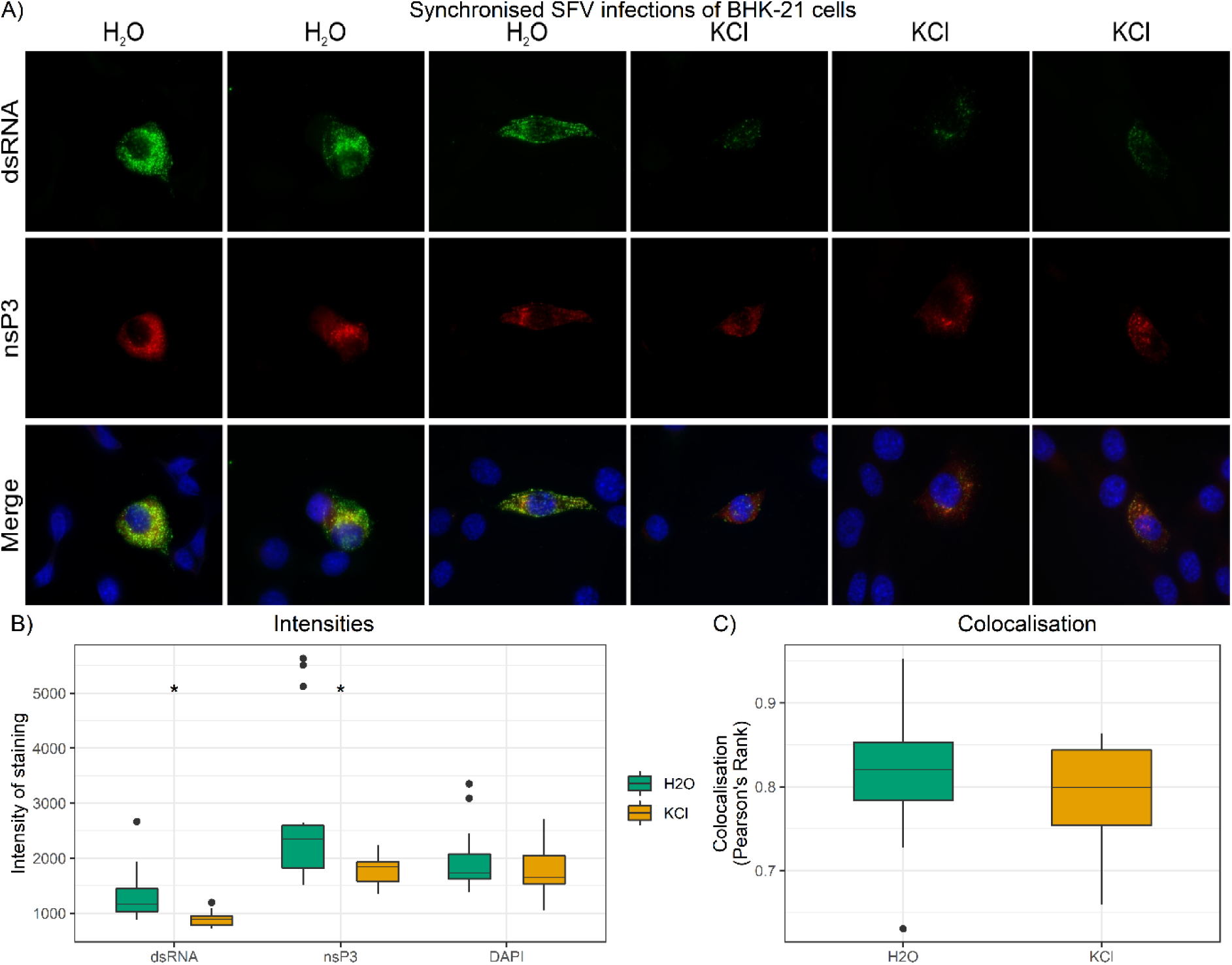
Effect of KCl on nsP3 and dsRNA staining in infected BHK-21 cells. A) Three representative confocal images each for untreated (H_2_O) and treated (KCl) BHK-21 cells following synchronised infections (MOI = 1) and fixing at 4 hpi. Cells were co-stained with J2 dsRNA (green) and nsP3 (red) antibodies. B) Intensities of dsRNA, nsP3 and DAPI staining were measured using ImageJ. C) Co-localisation of nsP3 and dsRNA was measured using ImageJ. Differences in measurements between treated and untreated cells were compared by Student t-tests and significance annotated as in Figure 1.

**Figure 8:**
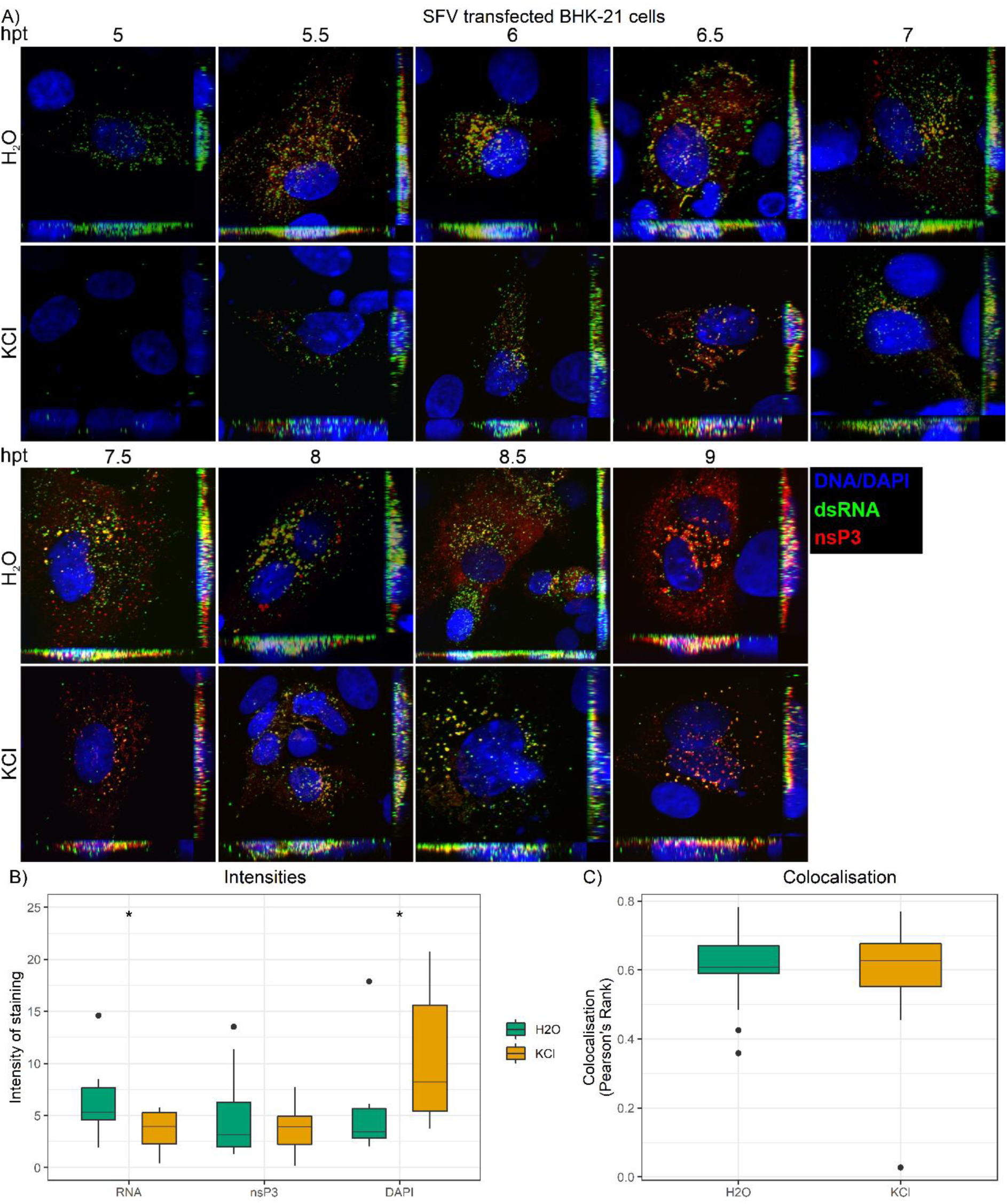
Effect of KCl on nsP3 and dsRNA staining in pCMV-SFV4 transfected cells. A) Representative images of BHK-21 cells transfected with pCMV-SFV4 in the presence or absence of KCl, fixed at 0.5 hour intervals between 5 and 9 hpt and then immunostained with J2 dsRNA (green) and anti-nsP3 (red) antibodies. B) Intensities were measured as in Figure 7. C) Colocalisation was measured as in Figure 7.

Taken together, the results of co-staining for nsP3 and dsRNA suggest KCl could be disrupting RC functionality or formation as these are the sites of viral RNA synthesis. Furthermore, several host proteins have been shown to associate with Alphavirus RCs [21, 24, 26, 57–60] so correct RC function appears to require assistance from cellular components. Co-localisation of nsP3 and dsRNA staining can be used to identify Alphavirus RCs where viral RNA synthesis is occurring [61, 62]. Analysis of co-localisation between these two viral components for synchronised infections (Figure 7C) and pCMV-SFV4 transfected (Figure 8C) BHK-21 cells showed there was no difference. Similar levels of co-localisation between nsP3 and dsRNA suggest the viral RNA being synthesised in KCl treated cells is still being synthesised in association with nsP3 at RCs.

### K+ channel inhibitors disrupt RNA replication and transcription

A transreplication assay was used to confirm the disruption of RNA synthesis by K^+^ channel inhibitors. BHK-21 cells were transfected with pWTnsP1234 (encodes nsP1234 from CMV promoter) or pMUTbsP1234 (encodes nsP1234 with mutation to prevent replicase activity) and pF4-Temp (FLuc under 5’ UTR promoter and GLuc under sub-genomic promoter) in the presence or absence of K^+^ channel inhibitors. Following subtraction of the background signal there was a significant reduction in FLuc and GLuc signal when assays were carried out in the presence of KCl or spermine compared to untreated controls (Figure 9A). There was no effect of the inhibitors on nsP3 abundance detected by western blot showing the inhibitors were not disrupting expression of nsP1234 from pWTnsP1234 (Figure 9B). The transreplication assay results show K^+^ channel inhibitors do not affect translation of nsP1234 from a CMV promoter, but they do disrupt RNA replication and transcription by the viral replicase.

**Figure 9:**
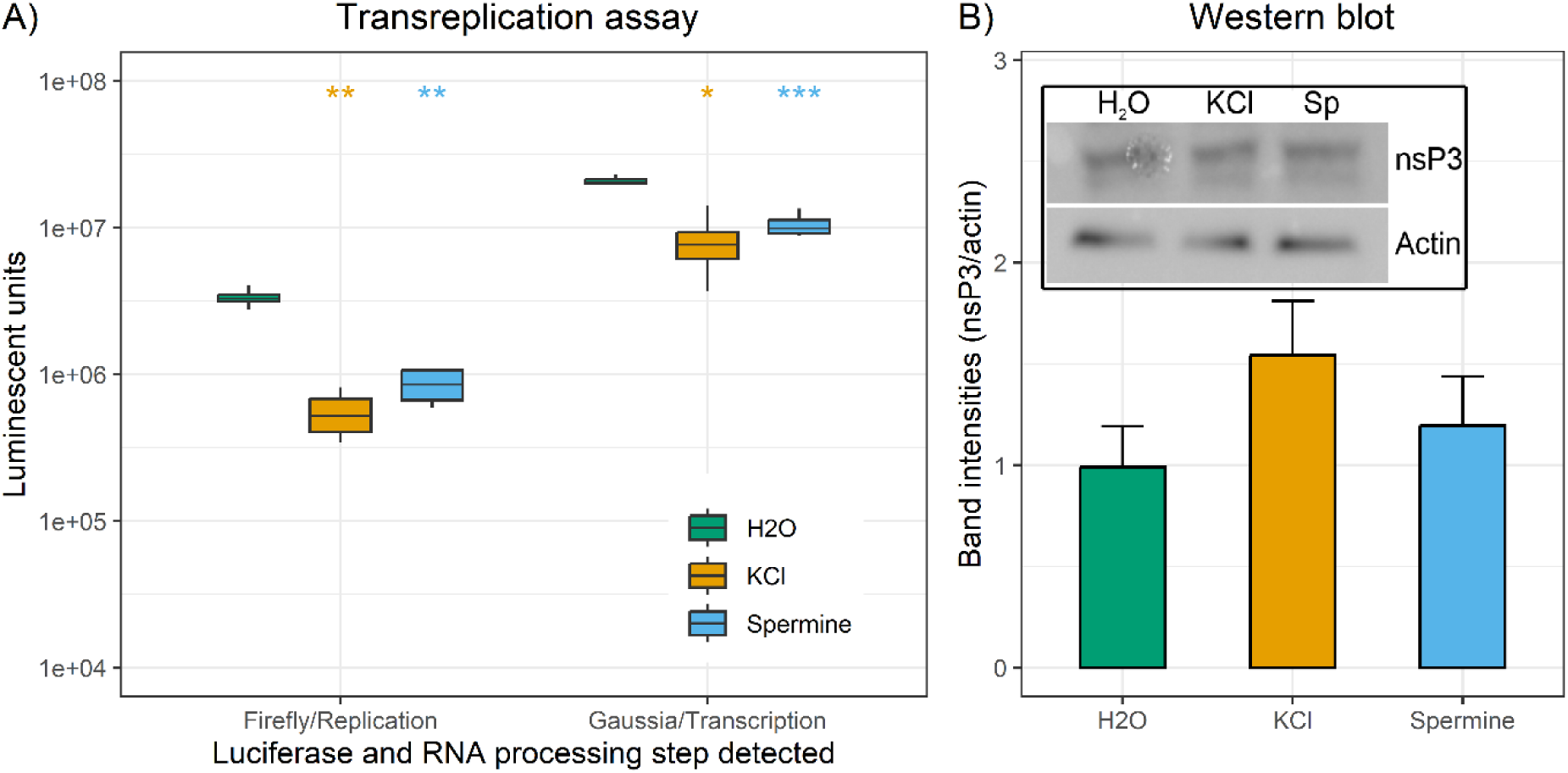
Transreplication assay shows K^+^ channel inhibitors disrupt SFV RNA replication and transcription. A) BHK-21 cells were transfected with pF4-Temp and pWTnsP1234 or pMUTnsP1234 in the presence or absence of the indicated inhibitor. FLuc and GLuc activity were measured at 18 hpt. Background luciferase activity (from pMUTnsP1234 transfected cells) was subtracted from luciferase activities for pWTnsP1234 experiments. B) Cell lysates underwent western blot using nsP3 and β-actin antibodies. Band intensities for nsP3 were normalised against β-actin band intensities. Significant results were annotated as in Figure 3.

## Discussion

### K^+^ channel inhibitors attenuate SFV replication

Ion channels are an attractive target for antivirals because they are involved in the efficient replication of some virus species and 13% of drugs currently in clinical use target ion channels so these drugs could be repurposed for treatment of viral diseases [33, 34]. Currently, no treatments exist to combat infections caused by members of the *Alphavirus* genus, which includes important human and animal pathogens. Therefore, the role of ion channels and their suitability as antiviral targets was investigated using a model *Alphavirus*, SFV. Initial screens showed multiple broad-range ion channel inhibitors could attenuate SFV replication or reduce SFV cytotoxicity so the therapeutic potential of targeting a range of ion channels for the treatment of *Alphavirus* infections could be pursued in the future. However, in this study K^+^ channels were investigated further because the only inhibitors to significantly attenuate SFV replication and reduce SFV cytotoxicity, were the K^+^ channel inhibitors KCl and TEA-Cl. The significant reduction in SFV titre up to at least 48 hpi in NIH 3T3 cells and up to at least 24 hpi in BHK-21 cells could be due to the intact IFN response in NIH 3T3 cells complimenting the antiviral effect of the inhibitors or it could indicate increased effectiveness of the K^+^ channel inhibitors in murine cells compared to hamster cells. KCl remained effective at disrupting SFV replication in *A. aegypti* cells suggesting that K^+^ channels or K^+^ ions where required for efficient replication in insect as well as mammalian cells.

Attenuation of SFV replication when KCl was administered at 2 or 4 hpi shows KCl can attenuate the virus when the replication cycle in underway. However, the failure to attenuate SFV infections at high MOI suggests KCl is not a potent inhibitor of SFV. The 2pK inhibitors, spermine and ruthenium red, were the only family-specific inhibitors to disrupt SFV replication suggesting 2pK channels are involved in SFV replication cycles. Normally, spermine and ruthenium red were slightly less effective than KCl so it cannot be ruled out that K^+^ channels targeted by KCl but not spermine and ruthenium red are also involved in the replication cycle. Spermine [52] and ruthenium red [55] both disrupt the 2pK channel, TASK-3, suggesting this could be one channel required for SFV replication.

### K^+^ channel inhibitors disrupt an early, post-entry step of SFV replication

K^+^ channel inhibitors remained effective when entry was bypassed and when they were administered after the 2-hour adsorption. Infecting cells in the presence of KCl also decreased the abundance of nsP3 and viral RNA at early timepoints post-infection. These results suggest K^+^ channel inhibitors disrupt an early, post-entry step of SFV replication. While KCl, spermine and ruthenium red were all effective when only added to infections post-adsorption, only the 2pK inhibitors caused a significant reduction in SFV titre when present only during the 2-hour adsorption. This suggests KCl does not affect entry of SFV but an effect of spermine and ruthenium red on entry cannot be ruled out. Alternatively, the results could suggest that spermine and ruthenium red have a stronger interaction with their cellular target so were not washed away during the PBS washes, or after being washed away, the effect of these inhibitors on K^+^ channels remains more long term.

### K^+^ channels inhibitors appear to disrupt SFV RNA synthesis

Having established KCl attenuates SFV replication at an early, post-entry step, attempts were made to identify the step of SFV replication targeted by these K^+^ channel inhibitor more precisely. The effect of KCl on both nsP3 and viral RNA abundance poses a conundrum as to whether it is affecting nsP synthesis or viral RNA synthesis because RNA is required as a template to generate nsP while nsP1234 is required to generate more viral RNA. Both the confocal images, where a larger reduction in dsRNA staining compared to nsP3 staining was observed, and the results of the transreplication assay, where there was no effect on nsP3 abundance but still an effect on RNA replication and transcription, suggest viral RNA synthesis is being disrupted by the K^+^ channel inhibitors. There is no evidence to suggest K^+^ channel inhibitors are disrupting RCs because there was no difference in co-localisation of nsP3 and dsRNA staining of confocal images. Though this could just be a result of fewer RCs synthesising less RNA in KCl treated cells compared to untreated, which would not necessarily result in a lower ratio of co-localisation – just reduced levels of staining as were observed. Results of the transreplication assay showed K^+^ channel inhibitors disrupt viral RNA replication as regulated by the 5’ UTR of the genome and transcription from the subgenomic promoter. This could indicate a non-specific effect on viral RNA synthesis, or it could indicate a targeted disruption of negative strand RNA synthesis because this is the template for replication and transcription.

2pK channels could be present on the membrane of the large cytoplasmic vacuoles or the spherules where viral and cellular proteins required for viral RNA synthesis associated as RCs. The endocytic pathway is hijacked by alphaviruses to generate these large cytoplasmic vacuoles and there is some evidence showing 2pk channels can be endocytosed. 2pK channels are normally found on the plasma membrane where they facilitate background/leak K^+^ currents but their activity is regulated by endocytosis as it removes them from the plasma membrane reducing the leak current [63–65]. Furthermore, 2pK channels were shown to be required for efficient replication of members of the *Peribunyaviridae* family [49] and subsequent studies by the same group showed K^+^ ions facilitated fusion of the viral envelope with the endosomal membrane [40, 41]. The association between 2pK channels and endosomes, although not conclusive, does suggest 2pK channels could at least be localised close to the sites of SFV RNA synthesis and potentially involved in this step of the replication cycle.

## Conclusions

Experiments investigating the ability of ion channel inhibitors to attenuate SFV replication showed that infecting cells in the presence of broad-range K^+^ channel inhibitors reproducibly decreased the virus titres obtained and protected cells from cytotoxic effects. KCl was shown to disrupt an early, post-entry step of SFV replication. Screens with family-specific K^+^ channel inhibitors showed only the 2pK inhibitors remained effective and there was evidence they are required for a post-entry step, although an effect of spermine and ruthenium red on SFV entry could not be ruled out. Results of confocal imaging and a transreplication assay suggested K^+^ channel inhibitors disrupt SFV RNA synthesis, but the precise mechanism or aspect of viral RNA synthesis targeted by these inhibitors has not been elucidated.

## Acknowledgements

We acknowledge the UCD Conway Institute Imaging Core, especially Professor Dimitri Scholz, for their help with the confocal imaging used in this manuscript.

